# DeepDist: real-value inter-residue distance prediction with deep residual convolutional network

**DOI:** 10.1101/2020.03.17.995910

**Authors:** Tianqi Wu, Zhiye Guo, Jie Hou, Jianlin Cheng

**Affiliations:** Department of Electrical Engineering and Computer Science, University of Missouri, Columbia, MO 65211, USA; Department of Computer Science, Saint Louis University, St. Louis, MO 63103, USA

## Abstract

**Motivation:** Driven by deep learning techniques, inter-residue contact/distance prediction has been significantly improved and substantially enhanced *ab initio* protein structure prediction. Currently all the distance prediction methods classify inter-residue distances into multiple distance intervals (i.e. a multi-classification problem) instead of directly predicting real-value distances (i.e. a regression problem). The output of the former has to be converted into real-value distances in order to be used in tertiary structure prediction.

**Results:** To explore the potentials of predicting real-value inter-residue distances, we develop a multi-task deep learning distance predictor (DeepDist) based on new residual convolutional network architectures to simultaneously predict real-value inter-residue distances and classify them into multiple distance intervals. We demonstrate that predicting the real-value distance map and multi-class distance map at the same time performs better than predicting real-value distances alone, indicating their complementarity. On 43 CASP13 hard domains, the average mean square error (MSE) of DeepDist’s real-value distance predictions is 0.896 Å when filtering out the predicted distance >=16 Å, which is lower than 1.003 Å of DeepDist’s multi-class distance predictions. When the predicted real-value distances are converted to binary contact predictions at 8Å threshold, the precisions of top L/5 and L/2 contact predictions are 78.6% and 64.5%, respectively, higher than the best results reported in the CASP13 experiment. These results demonstrate that the real-value distance prediction can predict inter-residue distances well and improve binary contact prediction over the existing state-of-the-art methods. Moreover, the predicted real-value distances can be directly used to reconstruct protein tertiary structures better than multi-class distance predictions due to the lower MSE.

## 1 Introduction

Recently, the accuracy of protein inter-residue contact prediction has been substantially increased due to the development of residue-residue co-evolution analysis methods effectively detecting the direct correlated mutations of contacted residues in the sequences of a protein family, such as Direct Coupling Analysis (DCA) (Weigt, et al., 2009), plmDCA (Ekeberg, et al., 2013), GREMLIN (Kamisetty, et al., 2013), CCMpred (Seemayer, et al., 2014), and PSICOV (Jones, et al., 2012). The capability of these methods to extract the correlated mutation information for contact prediction largely depends on the number of effective sequences in multiple sequence alignment (MSA) of a target protein. Due to the advancement in the DNA/RNA sequencing technology (Meyer, et al., 2008; Wilke, et al., 2016), many proteins have a lot of sufficiently diverse, homologous sequences that make their contact/distance prediction fairly accurate. However, for targets with a small number of effective homologous sequences (i.e. shallow sequence alignments), the co-evolutionary scores are noisy and not reliable for contact prediction. The problem can be largely addressed by using noisy co-evolutionary scores as input for advanced deep learning techniques that have strong pattern recognition power to predict inter-residue contacts and distances.

After deep learning contact prediction was first introduced for contact prediction in 2012 (Eickholt and Cheng, 2012; Di Lena, et al., 2012), different deep learning architectures have been designed to integrate traditional sequence features with inter-residue coevolution scores to substantially improve contact/distance prediction (Wang, et al., 2017; Adhikari, et al., 2018; Jones and Kandathil, 2018; Li, et al., 2019), even for some targets with shallow MSAs.

The improved contact predictions can be converted into inter-residue distance information, which has been successfully used with distance-based modeling methods such as CONFOLD (Adhikari, et al., 2015), CONFOLD2 (Adhikari and Cheng, 2018), and EVFOLD (Sheridan, et al., 2015) to build accurate tertiary structures for *ab initio* protein targets (Michel, et al., 2014; Monastyrskyy, et al., 2014).

In the most recent CASP13 experiment, several groups (e.g., AlphaFold (Senior, et al., 2020) and RaptorX (Xu, 2019)) applied deep learning techniques to classify inter-residue distances into multiple fine-grained distance intervals (i.e. predict the distance distribution) to further improve *ab initio* structure prediction substantially. However, the probabilities of a distance belonging to different intervals predicted by the multi-classification approach still need to be converted into a distance value to be used for tertiary structure modeling. There is a lack of regression methods to directly predict the exact real-value of inter-residue distances.

In this work, we develop a deep residual convolutional neural network method (DeepDist) to predict both the full-length real-value distance map and the multi-class distance map (i.e. distance distribution map) for a target protein. According to the test on 43 CASP13 hard domains (i.e. FM and FM/TBM domains; FM: free modeling; TBM: template-based modeling), 37 CASP12 hard (FM) domains, and 268 CAMEO targets, the method can predict inter-residue distance effectively and perform better than existing state-of-the-art methods in terms of the precision of binary contact prediction. We further show that predicting both real-value distance map and multi-class map simultaneously at the same time is more accurate than only predicting real-value distance map, demonstrating that the two kinds of predictions are complementary. They can be used together in multi-task learning to improve protein distance prediction.

## 2 Materials and Methods

### 2.1 Overview

The overall workflow of DeepDist is shown in **Fig. 1**. We use four sets of 2D co-evolutionary and sequence-based features to train four deep residual convolutional neural network architectures respectively to predict the Euclidean distance between residues in a protein target. Three of four feature sets are mostly coevolution-based features, i.e. covariance matrix (COV) (Jones and Kandathil, 2018), precision matrix (PRE) (Li, et al., 2019), and pseudolikelihood maximization matrix (PLM) (Seemayer, et al., 2014)) calculated from multiple sequence alignments. Considering that coevolution-based features sometimes cannot provide sufficient information, particularly when targets have shallow alignments, the fourth set of sequence-based features (OTHER), such as the sequence profile generated by PSI-BLAST (Bhagwat and Aravind, 2007), and solvent accessibility from PSIPRED (Jones, 1999) is used. The output of DeepDist is a real-value L × L distance map and a multi-class distance map (L: the length of the target protein). The two types of distance maps are generated by two prediction branches. For each branch, the final output is produced by the ensemble of four deep network models (COV_Net, PLM_Net, PRE_Net, and OTHER_Net) named after their input feature sets (COV, PLM, PRE, and OTHER). For the prediction of the multi-class distance map, we discretize the inter-residue distances into 25 bins: 1 bin for distance < 4.5Å, 23 bins from 4.5Å to 16Å at interval size of 0.5Å and a final bin for all distances greater than or equal to 16Å. For the real-value distance map, we simply use the true distance map of the native structure as targets to train deep learning models without discretization. Because large distances are not useful and not predictable, we only predict inter-residue distances less than 16 Å by filtering out true distances >= 16 Å.

**Fig. 1.**
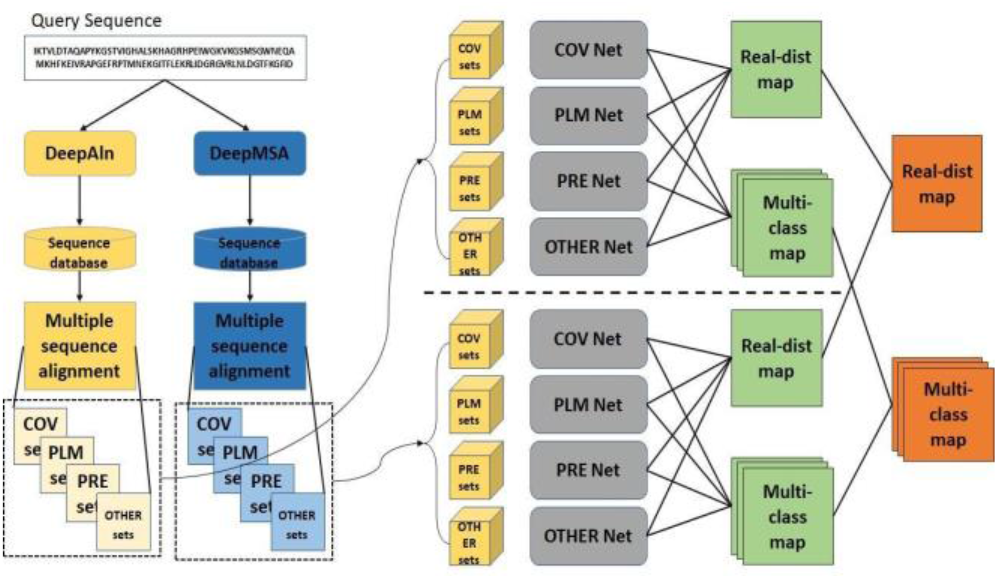
The overall workflow of DeepDist for both real-value distance map prediction and multi-class distance map prediction. Given a sequence, DeepAln and DeepMSA are called to search it against sequence databases to generate two kinds of multiple sequence alignments (MSAs), which are used to generate four sets of features (COV, PLM, PRE, OTHER), respectively. The four sets of features are used by four deep networks (COV Net, PLM Net, PRE Net, and OTHER Net) to predict both real-value distance (real-dist) map and multi-class distance (multi-class) map, respectively. The real-value distance maps (or multi-class distance maps) of the individual networks are averaged to produce the final real-value distance map (or multi-class distance map).

### 2.2 Datasets

We select targets from the training list used in DMPfold (Greener, et al., 2019) and extract their true structures from the Protein Data Bank (PDB) to create a training dataset. After filtering out the redundancy with the validation dataset and test datasets according 25% sequence identity threshold, 6463 targets are left in the training dataset. The validation set contains 144 targets used to validate DNCON2 (Adhikari, et al., 2018). The three blind test datasets are 37 CASP12 FM domains, 43 CASP13 FM and FM/TBM domains, and 268 CAMEO targets collected from 08/31/2018 to 08/24/2019.

### 2.3 Input Feature Generation

The sequence databases used to search for homologous sequences for feature generation include Uniclust30 (2017-10) (Mirdita, et al., 2017), Uniref90 (2018-04) and Metaclust50 (2018-01) (Steinegger and Söding, 2018), a customized database that combines Uniref100 (2018-04) and metagenomics sequence databases (2018-04), and NR90 database (2016). All of the sequence databases were constructed before the CASP13 experiment.

The three kinds of co-evolutionary features (i.e. COV, PRE, and PLM) are generated from multiple sequence alignment (MSA). Two methods, DeepMSA (Zhang, et al., 2019) and our in-house DeepAln, are used to generate MSA for a target. The outputs of both MSA generation methods are the combination of the iterative homologous sequence search of HHblits (Remmert, et al., 2012) and Jackhmmer (Eddy, 1992) on several sequence databases. The two methods differ in sequence databases used and the strategy of combining the output of HHblits and Jackhmmer searches. DeepMSA trims the sequence hits from Jackhmmer and performs sequence clustering, which shortens the time for constructing the HHblits database for the next round of search. To leverage its fast speed, we apply DeepMSA to search against a large customized sequence database that is composed of UniRef100 and metagenomic sequences. In contrast, DeepAln directly uses the full-length Jackhmmer hits for building HHblits customized databases and is slower. It is applied to the Metaclust sequences database. In addition to three kinds of co-evolutionary features, 2D features such as the coevolutionary contact scores generated by CCMpred, Shannon entropy sum, mean contact potential, normalized mutual information, and mutual information are also generated. Moreover, some other features used in DNCON2 including sequence profile, solvent accessibility, joint entropy, and Pearson correlation are also produced, which are collectively called OTHER feature.

The features above are generated for the MSAs of both DeepMSA and DeepAln. Each of them is used to train a deep model to predict both real-value distance map and multi-class distance map, resulting in 8 predicted real-value distance maps and 8 multi-class distance maps (**Fig. 1**).

### 2.4 Deep Network Architectures for Distance Prediction

We design different deep learning architectures that work well for four different types of input features, which are called COV_Net, PLM_Net, PRE_Net, and OTHER_Net (**Fig. 2**), respectively. PRE_Net and OTHER_Net share almost the same architecture with some minor difference.

**Fig. 2.**
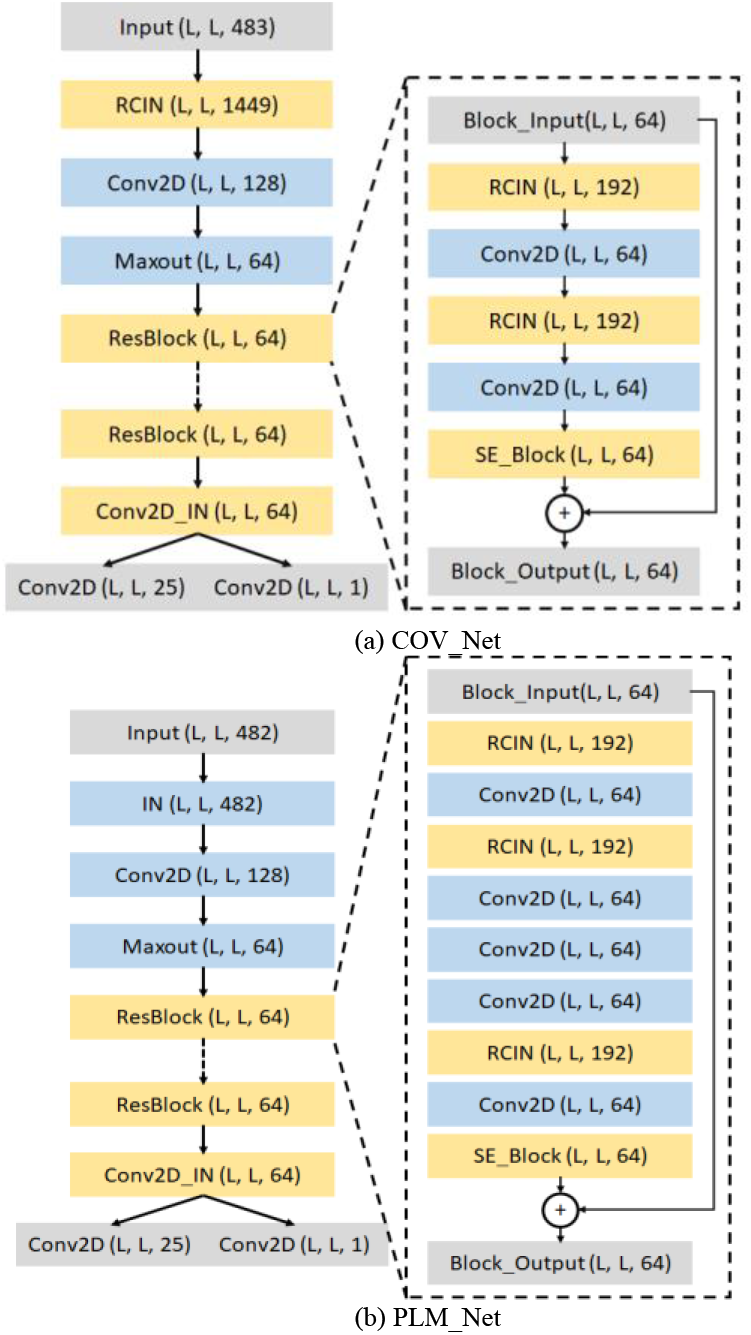
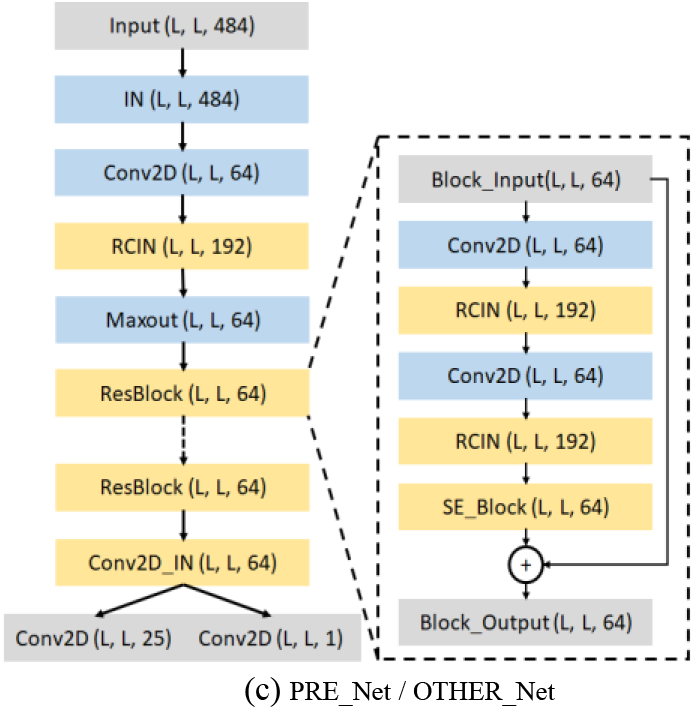
Deep network architectures for four deep residual network models (COV_Net, PRE_Net, PLM_Net, and OTHER_NET). RCIN: normalization layer; SE_block: squeeze- and-excitation block.

COV_Net (**Fig 2a**) uses as input the COV matrix along with sequence profile (PSSM), contact scores (CCMpred) and Pearson correlation. It starts with a normalization block called RCIN that contains instance normalization (IN) (Ulyanov, et al., 2016), row normalization (RN), column normalization (CN) (Mao, et al., 2019) and a ReLU (Nair and Hinton, 2010) activation function, followed by one convolutional layer with 128 kernels of size 1×1 and one Maxout (Goodfellow, et al., 2013) layer to reduce the input channel from 483 to 64. The output of Maxout is then fed into 16 residual blocks. Each residual block is composed of two RCIN normalization blocks, two convolutional layers that consist of 64 kernels of size 3×3 and one squeeze-and-excitation block (SE_block) (Hu, et al., 2018). The output feature maps from the block, together with the input of the block are added together as input for a ReLU activation function to generate the output of the residual block. The last residual block is followed by one convolutional instance normalization layer. The output of the layer is converted into two output maps simultaneously. One real-value distance map is obtained by a ReLU function through a convolution kernel of size 1×1, and one multi-class distance map with 25 output channels is obtained by a softmax function.

PLM_Net (**Fig. 2b**) uses as input the PLM matrix concatenated with the sequence profile (PSSM) and Pearson correlation. The input is first fed into an instance normalization layer, followed by one convolutional layer nand one Maxout layer. The output of Maxout is then fed into 20 residual blocks. Each residual block contains three RCIN blocks, four convolutional layers with 64 kernels of size 3×3, one SE_block and one dropout layer (Srivastava, et al., 2014) with a dropout rate of 0.2. The residual block is similar to the bottleneck residual block, except that the middle convolutional layer of kernel size 3×3 is replaced with three convolutional layers of kernel size 3×3, 7×1, 1×7, separately. The last residual block is followed by the same layers as in COV_Net to predict a real-value distance map and a multi-class distance map.

PRE_Net (**Fig. 2c**) uses as input the PRE matrix as well as entropy scores (joint entropy, Shannon entropy) and sequence profile (PSSM). An instance normalization layer is first applied to the input. Unlike COV_Net and PLM_Net, one convolutional layer with 64 kernels of size 1×1 and an RCIN block are applied after the instance normalization layer for dimensionality reduction. The output of the RCIN block is then fed through 16 residual blocks. Each residual block is made of two stacked sub-blocks (each containing one convolutional layer with 64 kernels of size 3×3, a RCIN block, a dropout layer with a dropout rate of 0.2, a SE_block, and the shortcut connection). The final output layers after the residual blocks are the same as in COV_Net.

OTHER_Net uses OTHER features as input. Its architecture is basically the same as PRE_Net, except that it has 22 residual blocks and there is no dropout layer in each residual block.

The final output of DeepDist is an average real-value distance map and an average multi-class distance map calculated from the output of the four individual network models, i.e. the output of the ensemble of the individual networks.

### 2.5 Training

The dimension of the input of COV_Net, PLM_Net, and PLM_Net is L×L×483, L×L×482, and L×L×484 respectively, which is very large and consumes a lot of memory. Therefore, we use data generators from Keras to load large feature data batch by batch. The batch size is set as 1. A normal initializer (He, et al., 2015) is used to initialize the network. For epochs ≤ 30, Adam optimizer (Kingma and Ba, 2014) is performed with an initial learning rate of 0.001. For epochs > 30, stochastic gradient descent (SGD) with momentum (Qian, 1999) is used instead, with the initial learning rate of 0.01 and the momentum of 0.9. The real-value distance prediction and multi-class distance classification are trained in two parallel branches. The mean squared error (MSE) and cross-entropy are used as their loss function, respectively. At each epoch, the precision of top L/2 long-range contact predictions derived from the average of the two contact maps converted from the real-value distance map and the multi-class distance map on the validation dataset is calculated. The inter-residue real-value distance map is converted to the contact map by inversing the predicted distance to obtain a relative contact probability (i.e. d_ij_: predicted distance between residues i and j; 1/d_ij_: relative contact probability score). The multi-class distance map is converted to the contact map by summing up the predicted probabilities of all the distance intervals <= 8Å as contact probabilities.

### 2.6 Ab Initio Protein Folding by Predicted Distances

We use distances predicted by DeepDist with our in-house tool – DFOLD (unpublished) built on top of CNS (Brünger, et al., 1998), a software package that implements distance geometry algorithm for NMR based structure determination, to construct 3D structure models. For the predicted real-value distance map, we select the predicted distances <= 15Å and with sequence separation >= 3 to generate the distance restraints between Cb-Cb atoms of residue pairs. 0.1Å is added to or subtracted from the predicted distances to set the upper and lower distance bounds. For the predicted multi-class distance map, we first convert the distance probability distribution matrix to a real-value distance map by setting each distance as the probability-weighted mean distance of all intervals for a residue pair and using the standard deviation to calculate the upper and lower distance bounds. Given a final real-value distance map, we prepare five different subsets of input distance constraints by filtering out distances ≥ *x* respectively, where *x* = 11Å, 12Å, 13Å, 14Å and 15Å. For each subset of distance constraints, we run DFOLD for 3 iterations. For each iteration, we generate 50 models and select the top five models ranked by the CNS energy score - the sum of all violations of all distance restraints used to generate a model. The top selected models generated from five subsets are further ranked by SBROD (Karasikov, et al., 2019). The final top one model is the one with the highest SBROD score. PSIPRED is used to predict the secondary structure to generate hydrogen bonds and torsion angle constraints for DFOLD to use.

## 3 Results

### 3.1 Comparing DeepDist with state-of-the-art methods on CASP12 and CASP13 datasets in terms of precision of binary contact predictions

As a multi-task predictor, our distance predictor DeepDist can not only classify each residue pair into distance intervals (multi-classification), but also predict its real-value distance (regression). We convert the predicted distances into contact maps in order to compare DeepDist with existing methods using the most widely used evaluation metrics – the precision of top L/5, L/2, L long-range contact predictions (sequence separation >= 24). **Fig. 3** reports the contact prediction precision of the multi-class distance prediction and the real-value distance prediction of DeepDist and several state-of-the art methods on two CASP test datasets (43 CASP13 FM and FM/TBM domains and 37 CASP12 FM domains). On the CASP13 dataset (**Fig 3a**), the precision of DeepDist is higher than the accuracy of three top methods (RaptorX-Contact (Xu and Wang, 2019), AlphaFold (Senior, et al., 2020), and TripletRes (Li, et al., 2019)) reported in CASP13 in almost all cases. For instance, the precision of top L/5 predicted contacts for DeepDist(multi-class) and DeepDist(real_dist) is 0.793 and 0.786 on the CASP13 dataset, respectively, higher than 0.744 of RaptorX-Contact. The multi-class distance prediction (DeepDist(multi-class)) works slightly better than the real-value distance prediction (DeepDist(real_dist)) according to this metric.

**Fig. 3.**
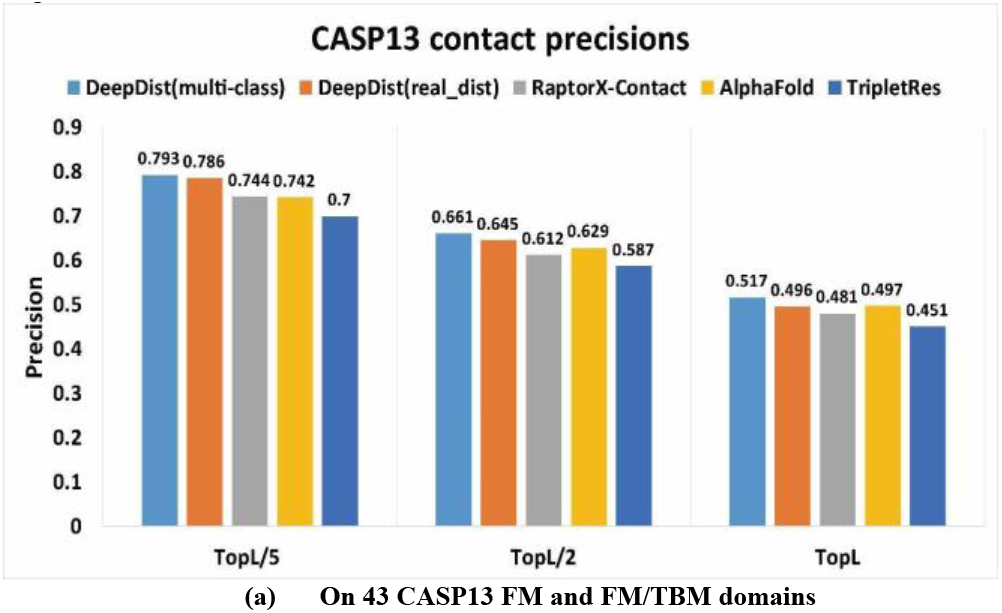
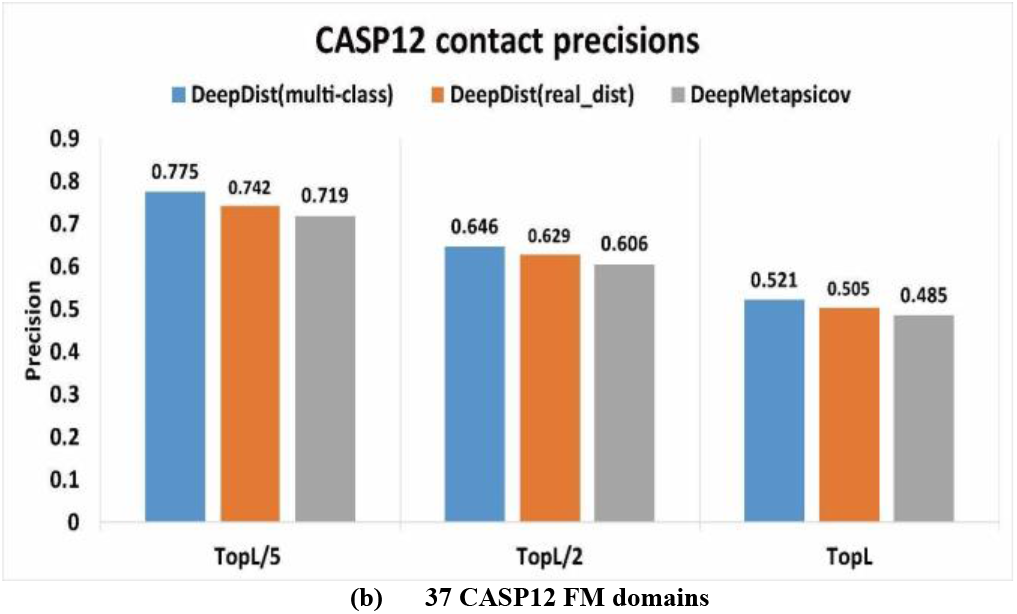
**(a)** Long-range contact prediction precision of DeepDist, RaptorX-Contact, AlphaFold,, TripletRes on CASP13 hard targets. “Top L/5”, “Top L/2” and “Top L” stands for the top L/5, L/2 and L predicted contacts, where L is the length of the domain. **(b)** Long-range contact prediction precision of DeepDist and DeepMetaPSICOV on CASP12 hard targets.

We also compare DeepDist with DeepMetaPSICOV on 37 CASP12 FM domains. To rigorously evaluate them, we ran DeepMetaPSICOV with the same sequence-based features (sequence profile from PSI-BLAST and solvent accessibility from PSIPRED) and MSAs used with DeepDist. Both multi-class distance prediction and real-value distance prediction of DeepDist perform consistently better than DeepMetaPSICOV (Kandathil, et al., 2019) (**Fig. 3b**).

### 3.2 Comparison of predicting real-value distance map and multi-class distance map simultaneously with predicting real-value distance map alone

In order to evaluate if predicting real-value distance map and multi-class distance map together improves the performance over predicting real-value distance map only, we apply the same deep learning architecture on the PLM input features in the two experimental settings. The precision of top L/5, top L/2, and L contact predictions, as well as MSE and Pearson correlation of predicted distances on 43 CASP13 FM and FM/TBM domains, are reported in **Table 1**. The results show that predicting the two at the same time is better than predicting real-value distances only, demonstrating coupling multi-class distance prediction with real-value distance prediction can improve the performance of real-value distance prediction according to all the metrics. DeepDist(real_dist) works slightly better in terms of MSE and Pearson’s correlation than DeepDist (multi-class), but slightly worse in terms of precision of top contact predictions.

**Table 1.** The results of predicting real-value distance map and multi-class distance map (DeepDist_PLM_Net (real-dist) and DeepDist_PLM_Net (multi-class) at the same time versus predicting real-value distance separately (real-dist ony) on 43 CASP13 hard domains. MSE: average mean square error between predicted distances and true distances; Pearson coefficient: the Pearson’s correlation between predicted distance and true distance.

|  | L/5 | L/2 | L | MSE | Pearson |
| --- | --- | --- | --- | --- | --- |
|  | (Precision) | (Precision) | (Precision) |  | coefficient |
| DeepDist_PLM_Net (real-dist) | 0.699 | 0.580 | 0.446 | 1.151 | 0.979 |
| DeepDist_PLM_Net (multi-class) | 0.711 | 0.587 | 0.457 | 1.206 | 0.978 |
| Real-dist only | 0.687 | 0.558 | 0.430 | 1.282 | 0.978 |

### 3.3 Comparison of the ensemble model based on four kinds of input and a single model based on one input

**Table 2** reports the performance of DeepDist (an ensemble of multiple models trained on four kinds of input) on the CASP13 dataset. The accuracy of DeepDist’s real-value distance prediction (DeepDist(real-dist)) and multi-class distance prediction (DeepDist(multi-class)) in Table 2 is substantially higher than the accuracy of a single deep model (DeepDist_PLM_Net) trained on one kind of feature - PLM in Table 1.

**Table 2.**
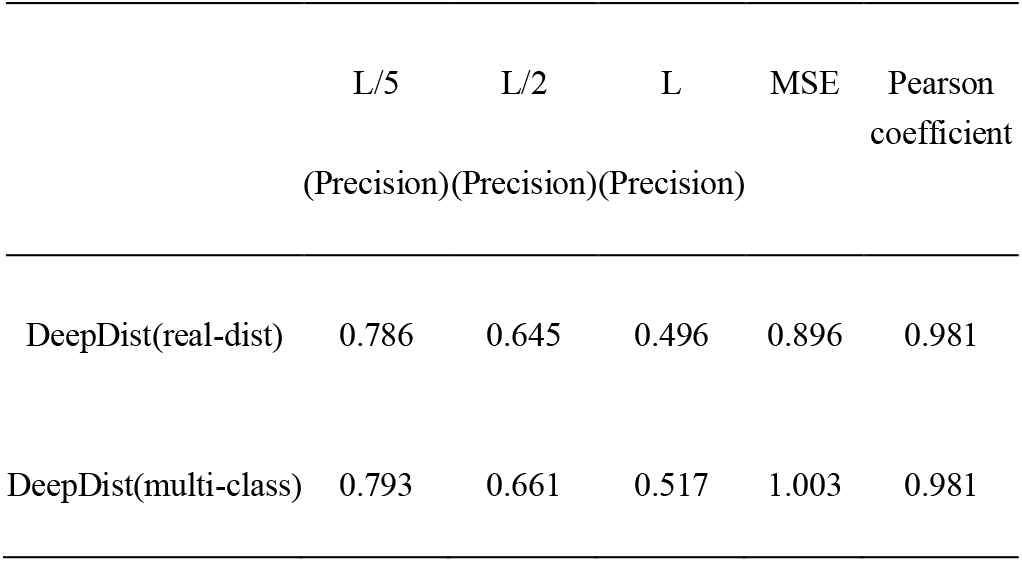
The performance of DeepDist on 43 CASP13 hard domains. DeepDist(real-dist): real-value distance prediction; DeepDist(multi-class): multi-class distance prediction

For instance, the precision for top L/5 contact prediction and MSE of DeepDist (real-dist) are 0.786 and 0.896 Å, better than 0.699 and 1.151 Å of DeepDist_PLM_Net (real-dist). The same results are observed for other single models trained on COV, PRE or OTHER features, separately. The results clearly demonstrate that the ensemble approach improves the accuracy of inter-residue distance prediction.

### 3.4 Comparison of real-value distance prediction and multi-class distance prediction of DeepDist in terms of contact prediction accuracy, MSE, and Pearson’s correlation

As shown in **Table 2**, the multi-class distance prediction of DeepDist is slightly better than the real-value distance prediction in terms of precision of contact prediction, but is a little worse in terms of MSE of predicted distance, and is the same in terms of Pearson’s correlation of predicted distance on the CASP13 dataset. Overall, their performance is comparable and the two kinds of predictions are complementary.

### 3.5 Comparison between real-value distance prediction and multi-class distance distribution prediction in term of 3D protein structure folding

We use the real-value distance map and multi-class distance map predicted by DeepDist with DFOLD to construct the 3D models for the 43 CASP13 hard domains respectively in order to compare their usefulness for 3D structure folding. **Table 3** shows the average TM-score of the top 1 model and the best model of the top 5 models of using real-value distances (DeepDist(real-dist)) and of using multi-class distances (DeepDist(multi-class)) on the 43 CASP13 FM and FM/TBM domains. The average TM-scores of top 1 and top 5 models generated from real-value distance predictions are 5.2%, and 3.1%, respectively, higher than those models generated from multi-class distance predictions.

**Table 3.**
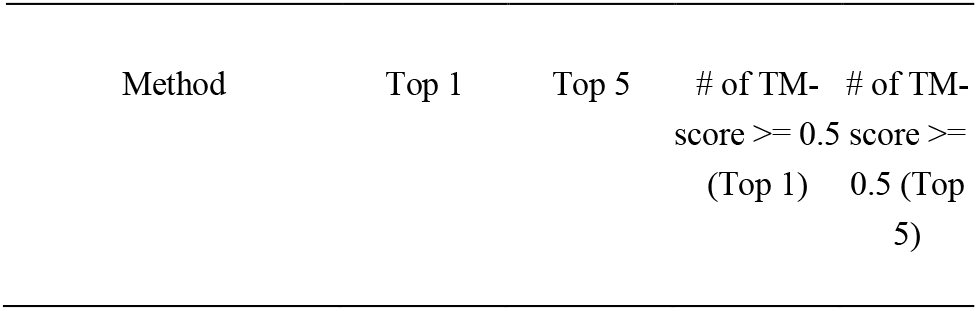
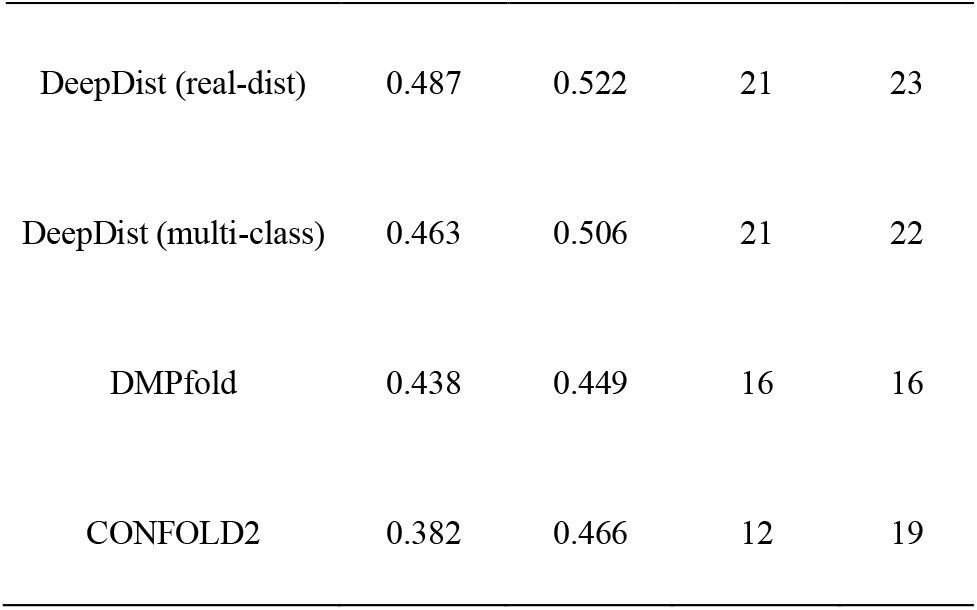
TM-scores of models on CASP13 43 FM and FM/TBM domains for four methods.

**Fig. 4** illustrates the distribution of TM-score of the top1 models of 43 CASP13 domains for DeepDist (real-dist) and DeepDist(multi-class). The distribution of DeepDist (real-dist) shift toward higher scores. The improvement of DeepDist (real-dist) over DeepDist(multi-class) is probably attributed to the reduction of MSE of predicted distances. The average MSE between the predicted real-value distance map and the true distance map is 0.8964 Å, which is lower than the average MSE (1.0037 Å) between the distance map converted from the predicted multi-class distance map and the true distance map.

**Fig. 4.**
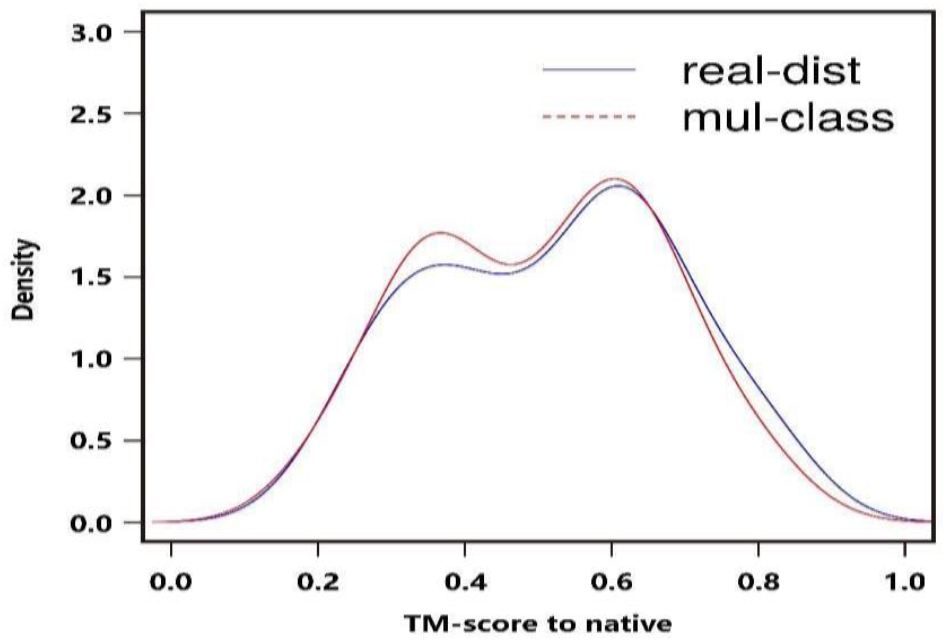
Distribution of TM-scores of the top 1 models of 43 CASP13 FM and FM/TBM domains, built from the real-value distance predictions and the multi-class distance predictions.

On the 43 CASP13 FM and FM/TBM domains, we also compared the models generated from the predicted distance of DeepDist with two popular ab initio distance-based model folding methods: DMPfold and CONFOLD2 (**Table 3**). For DMPfold, we applied the same sequence-based features and multiple sequence alignment used with DeepDist as input for DMPfold to build 3D models. For CONFOLD2, we converted the predicted distance map to the contact map as its input to build 3D models. As shown in **Table 3**, Both DeepDist and DMPfold have a much better performance than the contact-based method CONFOLD2, clearly demonstrating that the distance-based 3D modeling is better than contact-based 3D modeling. The average TM-score of DeepDist (real-dist) is 0.487, higher than 0.438 of DMPfold, probably due to more accurate distance prediction made by DeepDist. Considering top 5 models, DeepDist(real_dist) folds 23 out of 43 domains (TM-score > 0.5) correctly, higher than 16 of DMPfold. **Fig. 5** shows five high-quality CASP13 models built from the predicted real-value distances that have the TM-scores >= 0.7.

**Fig. 5.**
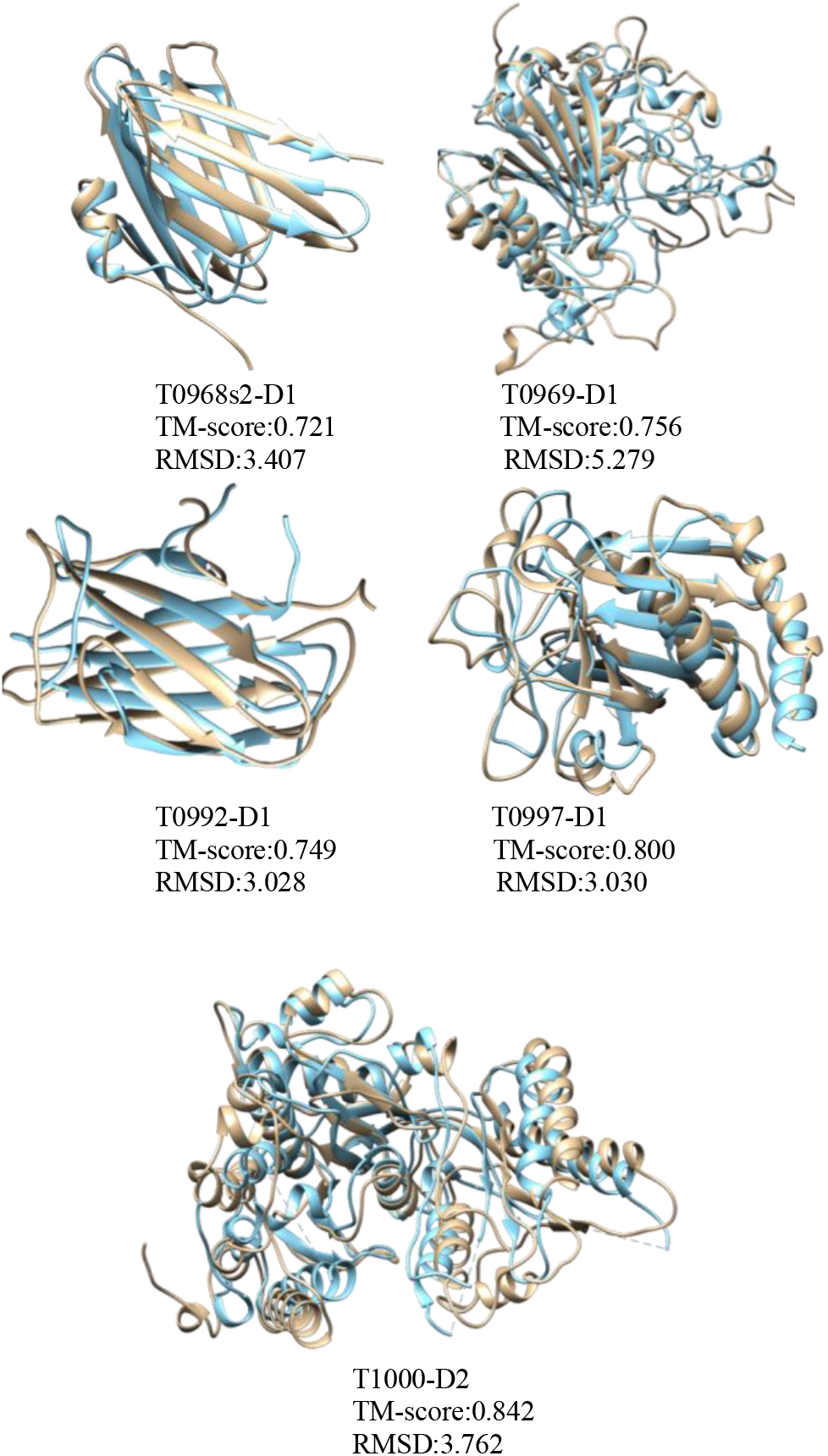
Five high-quality CASP13 models (TM-score >= 0.7) generated from DeepDist real-value distance predictions. Brown: model; Blue: native structure.

### 3.6 The relationship between 3D models reconstructed from predicted real-value distances and multiple sequence alignments

The main input features used with DeepDist are derived from MSAs. **Fig. 6** plots the TM-scores of top 1 models of 43 CASP13 domains against the natural logarithm of number of effective sequences in their MSAs. There is a moderate correlation (Pearson’s correlation = 0.66) between the two. Moreover, 3D models for 6 domains (T0957s2-D1, T0958-D1, T0986s2-D1, T0987-D1, T0989-D1, and T0990-D1) with shallow alignments (the number of effective sequences (Neff) in the alignment < 55) have TM-score > 0.5 (i.e. TM-score 0.568, 0.644, 0.658, 0.555, 0.545 and 0.593, respectively), indicating DeepDist works well on some targets with shallow alignments.

**Fig. 6.**
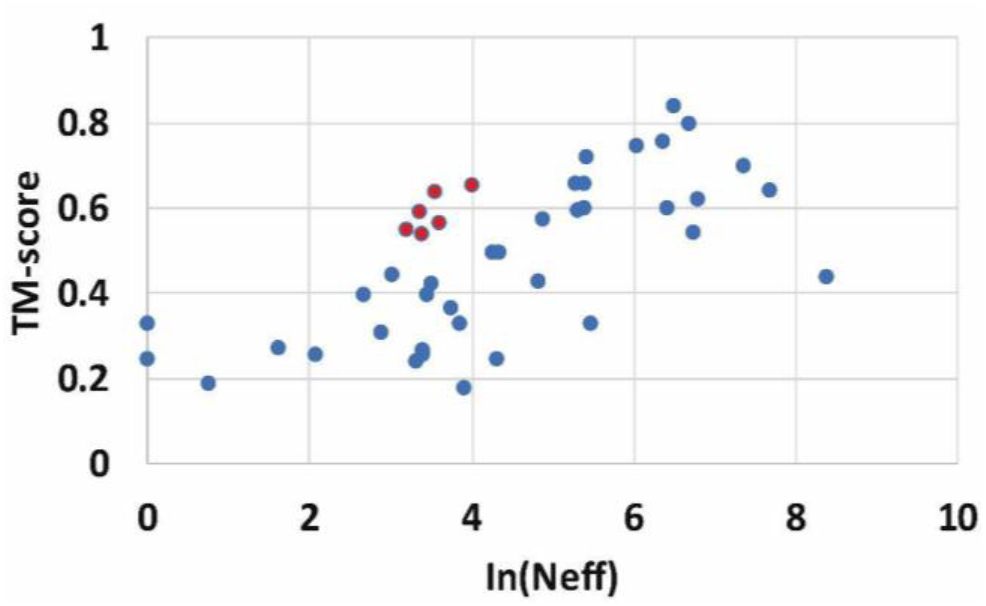
The quality of the top 1 models folded from DeepDist real-value distance predictions versus the logarithm of number of effective sequences (Neff) on 43 CASP13 FM and FM/TBM domains. The six points in red denote domains with Neff < 55 and TM-score > 0.5.

### 3.7 Evaluation on CAMEO targets

In order to further evaluate DeepDist on a large dataset, we test DeepDist on 268 CAMEO targets selected from 08/31/2018 to 08/24/2019. The average precision of the top L/5 or L/2 long-range inter-residue contact prediction converted from the real-value distance prediction is 0.691, and 0.598, respectively. 191 out of 268 targets have the long-range top L/5 contact prediction precision >= 0.7. **Fig. 7** shows 5 high-quality models constructed from DeepDist predicted real-value distances. For the 14 targets with the number of effective sequences less than or equal to 50, the average top L/5 and top L/2 long-range contact prediction precision is 0.696 and 0.515, which is reasonable. Using the predicted distance to build 3D structures for the 14 targets, five of them have models with TM-score > 0.5. This further confirms that DeepDist’s predicted distances can fold some proteins with very shallow alignments correctly.

**Fig. 7.**
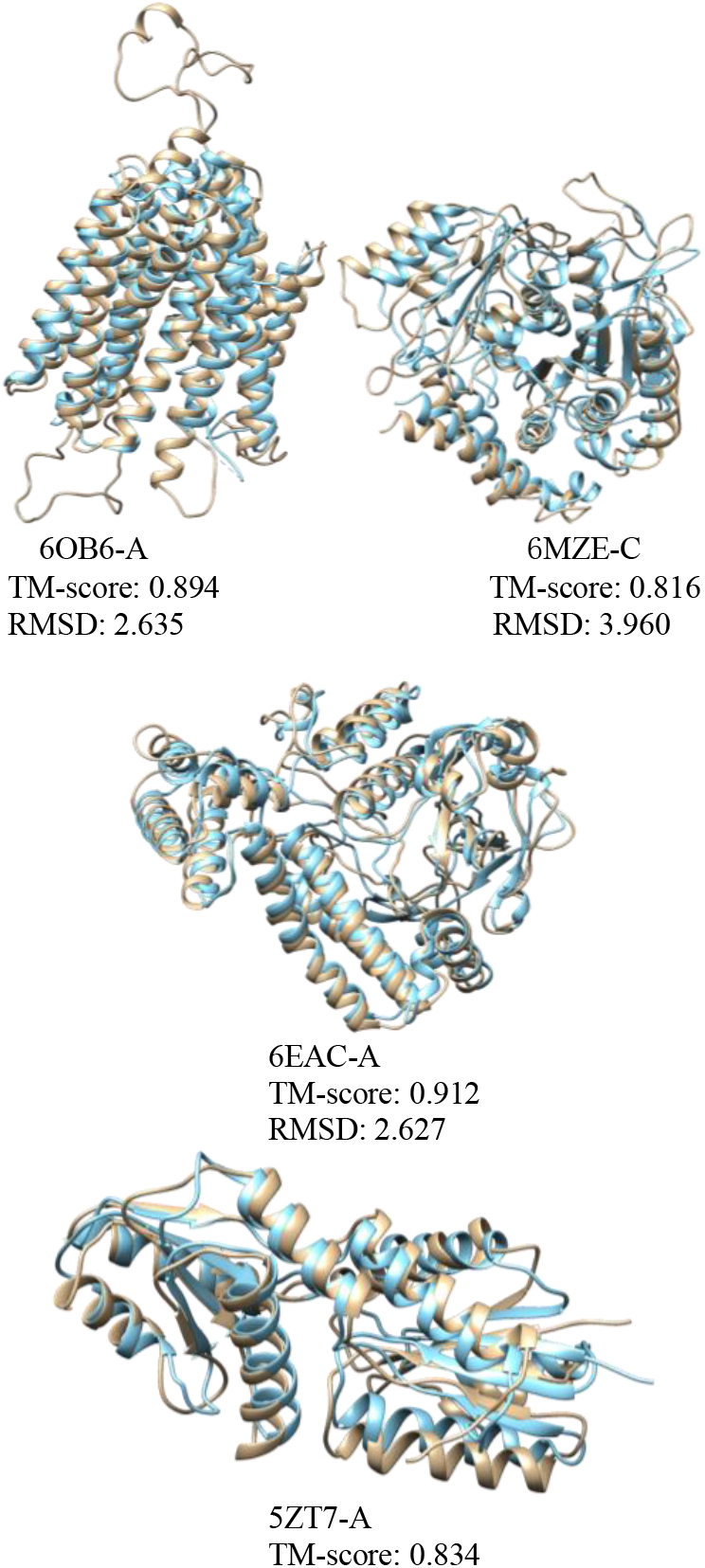
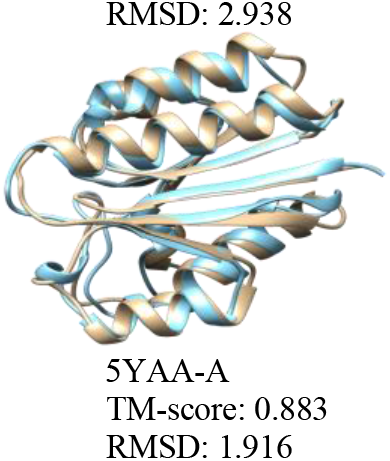
High-quality 3D models for five CAMEO targets constructed from DeepDist predicted real-value distances. The model is shown in brown and the native structure is shown in blue.

## Conclusion

We develop an inter-residue distance predictor DeepDist based on new deep residual convolutional neural networks to predict both real-value distance map and multi-class distance map simultaneously. We demonstrate that predicting the two at the same time yields higher accuracy in real-value distance prediction than predicting real-value distance alone. The overall performance of DeepDist’s real-value distance prediction and multi-class distance prediction is comparable according to multiple evaluation metrics. Both kinds of distance predictions of DeepDist are more accurate than the state-of-the-art methods on the CASP13 hard targets. Moreover, DeepDist can work well on some targets with shallow multiple sequence alignments. And the real-value distance predictions can be used to reconstruct 3D protein structures better than predicted multi-class distance predictions, showing that predicting real-value inter-residue distances can add the value on top of existing distance prediction approaches.

## References

Adhikari, B., et al. CONFOLD: residuer-esiduecontact-guidedab initio protein folding. Proteins: Structure, Function, and Bioinformatics 2015;83(8):1436–1449.

Adhikari, B. and Cheng, J. CONFOLD2: improved contact-driven ab initio protein structure modeling. BMC bioinformatics 2018;19(1):22.

Adhikari, B., Hou, J. and Cheng, J. DNCON2: improved protein contact prediction using two-level deep convolutional neural networks. Bioinformatics 2018;34(9):1466–1472.

Bhagwat, M. and Aravind, L. Psi-blast tutorial. In, Comparative genomics. Springer; 2007. p. 177–186.

Brünger, A.T., et al. Crystallography & NMR system: A new software suite for macromolecular structure determination. Acta Crystallographica Section D: Biological Crystallography 1998;54(5):905–921.

Di Lena, P., Nagata, K. and Baldi, P. Deep architectures for protein contact map prediction. Bioinformatics 2012;28(19):2449–2457.

Eddy, S. HMMER uesr’s gudie. Department of Genetics, Washington University School of Medicine 1992;2(1):13.

Eickholt, J. and Cheng, J. Predicting protein residue-residue contacts using deep networks and boosting. Bioinformatics 2012;28(23):3066–3072.

Ekeberg, M., et al. Improved contact prediction in proteins: using pseudolikelihoods to infer Potts models. Physical Review E 2013;87(1):012707.

Goodfellow, I.J., et al. Maxout networks. arXiv preprint arXiv:1302.4389 2013.

Greener, J.G., Kandathil, S.M. and Jones, D.T. Deep learning extends de novo protein modelling coverage of genomes using iteratively predicted structural constraints. Nature communications 2019;10(1):1–13.

He, K., et al. Delving deep into rectifiers: Surpassing human-level performance on imagenet classification. In, Proceedings of the IEEE international conference on computer vision. 2015. p. 1026–1034.

Hu, J., Shen, L. and Sun, G. Squeeze-and-excitation networks. In, Proceedings of the IEEE conference on computer vision and pattern recognition. 2018. p. 7132–7141.

Jones, D.T. Protein secondary structure prediction based on position-specific scoring matrices. Journal of molecular biology 1999;292(2):195–202.

Jones, D.T., et al. PSICOV: precise structural contact prediction using sparse inverse covariance estimation on large multiple sequence alignments. Bioinformatics 2012;28(2):184–190.

Jones, D.T. and Kandathil, S.M. High precision in protein contact prediction using fully convolutional neural networks and minimal sequence features. Bioinformatics 2018;34(19):3308–3315.

Kamisetty, H., Ovchinnikov, S. and Baker, D. Assessing the utility of coevolution-based residue-residue contact predictions in a sequence- and structure-rich era. Proceedings of the National Academy of Sciences 2013;110(39):15674–15679.

Kandathil, S.M., Greener, J.G. and Jones, D.T. Prediction of interresidue contacts with DeepMetaPSICOV in CASP13. Proteins: Structure, Function, and Bioinformatics 2019;87(12):1092–1099.

Karasikov, M., Pagès, G. and Grudinin, S. Smooth orientation-dependent scoring function for coarse-grained protein quality assessment. Bioinformatics 2019;35(16):2801–2808.

Kingma, D.P. and Ba, J. Adam: A method for stochastic optimization. arXiv preprint arXiv:1412.6980 2014.

Li, Y., et al. ResPRE: high-accuracy protein contact prediction by coupling precision matrix with deep residual neural networks. Bioinformatics 2019;35(22):4647–4655.

Li, Y., et al. Ensembling multiple raw coevolutionary features with deep residual neural networks for contact-map prediction in CASP13. Proteins: Structure, Function, and Bioinformatics 2019;87(12):1082–1091.

Mao, W., et al. AmoebaContact and GDFold as a pipeline for rapid de novo protein structure prediction. Nature Machine Intelligence 2019:1–9.

Meyer, F., et al. The metagenomics RAST server-a public resource for the automatic phylogenetic and functional analysis of metagenomes. BMC bioinformatics 2008;9(1):386.

Michel, M., et al. PconsFold: improved contact predictions improve protein models. Bioinformatics 2014;30(17):i482–i488.

Mirdita, M., et al. Uniclust databases of clustered and deeply annotated protein sequences and alignments. Nucleic acids research 2017;45(D1):D170–D176.

Monastyrskyy, B., et al. Evaluation of residue–residue contact prediction in CASP10. Proteins: Structure, Function, and Bioinformatics 2014;82:138–153.

Nair, V. and Hinton, G.E. Rectified linear units improve restricted boltzmann machines. In, Proceedings of the 27th international conference on machine learning (ICML-10). 2010. p. 807–814.

Qian, N. On the momentum term in gradient descent learning algorithms. Neural networks 1999;12(1):145–151.

Remmert, M., et al. HHblits: lightning-fast iterative protein sequence searching by HMM-HMM alignment. Nature methods 2012;9(2):173.

Seemayer, S., Gruber, M. and Söding, J. CCMpred—fast and precise prediction of protein residue–residue contacts from correlated mutations. Bioinformatics 2014;30(21):3128–3130.

Senior, A.W., et al. Improved protein structure prediction using potentials from deep learning. Nature 2020:1–5.

Sheridan, R., et al. Evfold. org: Evolutionary couplings and protein 3d structure prediction. BioRxiv 2015:021022.

Srivastava, N., et al. Dropout: a simple way to prevent neural networks from overfitting. The journal of machine learning research 2014;15(1):1929–1958.

Steinegger, M. and Söding, J. Clustering huge protein sequence sets in linear time. Nature communications 2018;9(1):1–8.

Ulyanov, D., Vedaldi, A. and Lempitsky, V. Instance normalization: The missing ingredient for fast stylization. arXiv preprint arXiv:1607.08022 2016.

Wang, S., et al. Accurate de novo prediction of protein contact map by ultra-deep learning model. PLoS computational biology 2017;13(1):e1005324.

Weigt, M., et al. Identification of direct residue contacts in protein-protein interaction by message passing. Proceedings of the National Academy of Sciences 2009;106(1):67–72.

Wilke, A., et al. The MG-RAST metagenomics database and portal in 2015. Nucleic acids research 2016;44(D1):D590–D594.

Xu, J. Distance-based protein folding powered by deep learning. Proceedings of the National Academy of Sciences 2019;116(34):16856–16865.

Xu, J. and Wang, S. Analysis of distance-based protein structure prediction by deep learning in CASP13. Proteins: Structure, Function, and Bioinformatics 2019;87(12):1069–1081.

Zhang, C., et al. DeepMSA: constructing deep multiple sequence alignment to improve contact prediction and fold-recognition for distant-homology proteins. Bioinformatics 2019.

